# Why Does the Neocortex Have Columns, A Theory of Learning the Structure of the World

**DOI:** 10.1101/162263

**Authors:** Jeff Hawkins, Subutai Ahmad, Yuwei Cui

## Abstract

Neocortical regions are organized into columns and layers. Connections between layers run mostly perpendicular to the surface suggesting a columnar functional organization. Some layers have long-range excitatory lateral connections suggesting interactions between columns. Similar patterns of connectivity exist in all regions but their exact role remain a mystery. In this paper, we propose a network model composed of columns and layers that performs robust object learning and recognition. Each column integrates its changing input over time to learn complete predictive models of observed objects. Excitatory lateral connections across columns allow the network to more rapidly infer objects based on the partial knowledge of adjacent columns. Because columns integrate input over time and space, the network learns models of complex objects that extend well beyond the receptive field of individual cells. Our network model introduces a new feature to cortical columns. We propose that a representation of location relative to the object being sensed is calculated within the sub-granular layers of each column. The location signal is provided as an input to the network, where it is combined with sensory data. Our model contains two layers and one or more columns. Simulations show that using Hebbian-like learning rules small single-column networks can learn to recognize hundreds of objects, with each object containing tens of features. Multi-column networks recognize objects with significantly fewer movements of the sensory receptors. Given the ubiquity of columnar and laminar connectivity patterns throughout the neocortex, we propose that columns and regions have more powerful recognition and modeling capabilities than previously assumed.

## INTRODUCTION

The neocortex is complex. Within its 2.5mm thickness are dozens of cell types, numerous layers, and intricate connectivity patterns. The connections between cells suggest a columnar flow of information across layers as well as a laminar flow within some layers. Fortunately, this complex circuitry is remarkably preserved in all regions, suggesting that a canonical circuit consisting of columns and layers underlies everything the neocortex does. Understanding the function of the canonical circuit is a key goal of neuroscience.

Over the past century, several theories have been proposed to explain the existence of cortical layers and columns. One theory suggested these anatomical constructs minimize the amount of wiring in cortical tissues (Shipp et al., 2007). Some researchers suggested there should be functional differentiation of different cortical layers that match the anatomical structure (Douglas and Martin, 2004). Others have proposed that long-range laminar connections contribute to attention-related changes in receptive field properties (Raizada and Grossberg, 2003). Recent advances in recording technologies now enable detailed recording of activity in the micro-circuitry of cortical columns. However, despite these advances, the function of networks of neurons organized in layers and columns remains unclear, and assigning any function to columns remains controversial (Horton and Adams, 2005).

Lacking a theory of why the neocortex is organized in columns and layers, almost all artificial neural networks, such as those used in deep learning (LeCun et al., 2015) and spiking neural networks (Maass, 1997), do not include these features, introducing the possibility they may be missing key functional aspects of biological neural tissue. To build systems that work on the same principles as the neocortex we need an understanding of the functional role of columnar and laminar projections.

Cellular layers vary in the connections they make, but a few general rules have been observed. Cells in layers that receive direct feedforward input do not send their axons outside the local region and they do not form long distance horizontal connections within their own layer. Cells in layers that are driven by input layers form long range excitatory connections within their layer, and also send an axonal branch outside of the region, constituting an output of the region. This two-layer input-output circuit is a persistent feature of cortical regions. The most commonly recognized instance involves layer 4 and Layer 2/3. Layer 4 receives feedforward input. It projects to layer 2/3 which is an output layer (Douglas and Martin, 2004; Shipp et al., 2007). Upper layer 6 also receives feedforward input (Thomson, 2010). It projects to layer 5, which is an output layer (Douglas and Martin, 2004; Guillery and Sherman, 2011), and therefore layers 6 and 5 may be a second instance of the two-layer input-output circuit. The prevalence of this two-layer connection motif suggests it plays an essential role in cortical processing.

In this paper, we introduce a theory of how columns and layers learn the structure of objects in the world. It is a sensorimotor theory in that learning and inference require movement of sensors relative to objects. We also introduce a network model based on the theory. The network consists of one or more columns, where each column contains an input layer and an output layer. First, we show how even a single column can learn the structure of complex objects. A single column can only sense a part of an object at any point in time, however, the column will be exposed to multiple parts of an object as the corresponding sensory organ moves. While the activation in the input layer changes with each movement of the sensor, the activation in the output layer remains stable, associating a single output representation with a set of feature representations in the input layer. Thus, a single cortical column can learn models of complete objects through movement. These objects can be far larger than any individual cell’s receptive field.

Next, we show how multiple columns collaborate via long-range intralaminar connections. At any point in time, each column has only partial knowledge of the object it is observing, yet adjacent columns are typically sensing the same object, albeit at different locations on the object. Long range excitatory connections in the output layer allow multiple columns to rapidly reach a consensus of what object is being observed. Although learning always requires multiple sensations via movement, inference with multiple columns can often occur in a single or just a few sensations. Through simulation we illustrate that our model can learn the structure of complex objects, it infers quickly, and it has high capacity.

A key component of our theory is the presence in each column of a signal representing location. The location signal represents an “allocentric” location, meaning it is a location relative to the object being sensed. In our theory, the input layer receives both a sensory signal and the location signal. Thus, the input layer knows both what feature it is sensing and where the sensory feature is on the object being sensed. The output layer learns complete models of objects as a set of features at locations. This is analogous to how computer-aided-design programs represent multi-dimensional objects.

Because different parts of a sensory array (for example different fingers or different parts of the retina) sense different parts of an object, the location signal must be calculated uniquely for each sensory patch and corresponding area of neocortex. We propose that the location signal is calculated in the sub-granular layers of cortical columns and is passed to input layer 4 via projections from layer 6.

It is important to note that we deduced the existence of the allocentric location signal. We first deduced its presence by considering how fingers can predict what they will sense while moving and touching an object. However, we believe the location signal is present in all neocortical regions. We show empirical evidence in support of this hypothesis. Although we cannot yet propose a complete mechanism for how the location signal is derived, the task of determining location and predicting new locations based on movement is similar to what grid cells do in the medial entorhinal cortex. Grid cells offer an existence proof that predictive models of allocentric location are possible, and they suggest mechanisms for how the location signal might be derived in cortical columns.

The theory is consistent with a large body of anatomical and physiological evidence. We discuss this support and propose several predictions that can be used to further test the theory.

## MODEL

### Motivation

Our research is focused on how networks of neurons in the neocortex learn predictive models of the world. Previously, we introduced a network (Hawkins and Ahmad, 2016) that learns a predictive model of naturally changing sensory sequences. In the present paper, we extend this network to address the related question of how the neocortex learns a predictive model of static objects, where the sensory input changes due to our own movement.

A simple thought experiment may be useful to understand our model. Imagine you reach your hand into a black box and try to determine what object is in the box, say a coffee cup. Using only one finger it is unlikely you could identify the object with a single touch. However, after making one contact with the cup, you move your finger and touch another location, and then another. After a few touches, you identify the object as a coffee cup. Recognizing the cup requires more than just the tactile sensation from the finger, the brain must also integrate knowledge of how the finger is moving, and hence where it is relative to the cup. Once you recognize the cup, each additional movement of the finger generates a prediction of where the finger will be on the cup after the movement, and what the finger will feel when it arrives at the new location. This is the first problem we wanted to address, how a small sensory array (e.g. the tip of a finger) can learn a predictive model of three dimensional objects by integrating sensation and movement-derived location information.

If you use two fingers at a time you can identify the cup with fewer movements. If you use five fingers you will often be able to identify an object with a single grasp. This is the second problem we wanted to address, how a set of sensory arrays (e.g. tips of multiple fingers) work together to recognize an object faster than they can individually.

Somatic inference is obviously a sensorimotor problem. However, vision and audition are also sensorimotor tasks. Therefore, the mechanisms underlying sensorimotor learning and inference should exist in all sensory regions, and any proposed network model should map to the detailed anatomical and physiological properties that exist in all cortical regions. This mapping, an explanation of common cortical circuitry, is a third goal of our model.

### Model description

Our model extends previous work showing how a single layer of pyramidal neurons can learn sequences and make predictions (Hawkins and Ahmad, 2016). The current model consists of two layers of pyramidal neurons arranged in a column. The model has one or more of these columns (**Figure. 1A**). Each cortical column processes a subset of the sensory input space and is exposed to different parts of the world as the sensors move. The goal is to have the output layer of each column converge on an object representation that is consistent with the accumulated sensations over time and across all columns.

The input layer of each column in our model receives a sensory input and a location input. The sensory input is a sparse binary array representing the current feature in its input space. The location input is a sparse binary array representing the location of the feature on the object. There are numerous observations in the neocortex that receptive fields are modified by location information. Grid cells in the entorhinal cortex also solve a similar location encoding problem and therefore represent a model of how location might be derived in the neocortex. We explore these ideas further in the discussion section. For our model we require a) that the location of a feature on an object is independent of the orientation of the object, and b) that nearby locations have similar representations. The first property allows the system to make accurate predictions when the object is sensed in novel positions relative to the body. The second property enables noise tolerance – you don’t have to always sense the object in precisely the same locations.

**Figure 1.**
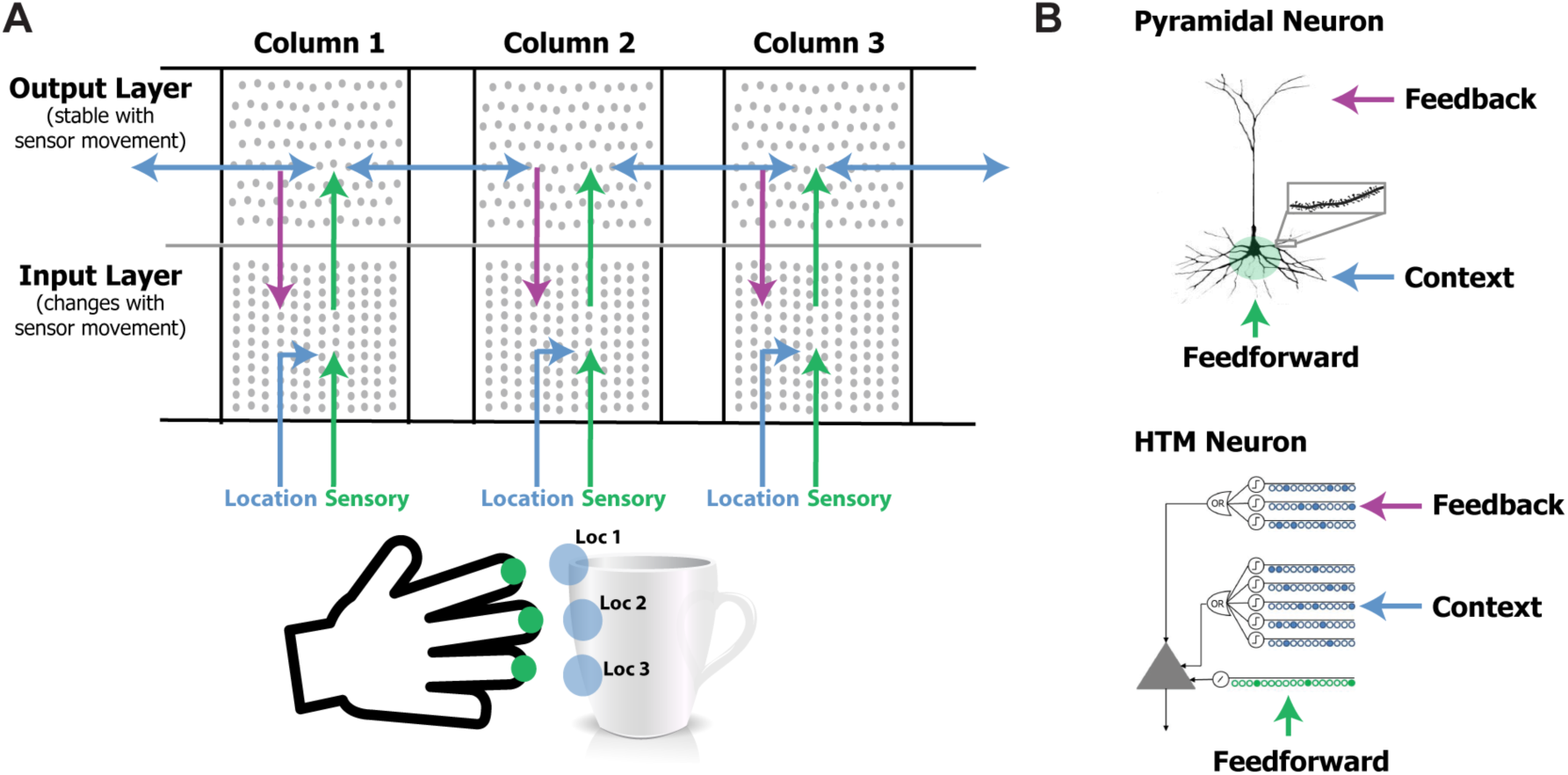
**A.** Our network model contains one or more laterally connected cortical columns (three shown). Each column receives feedforward sensory input from a different sensor array (e.g. different fingers or adjacent areas of the retina (not shown)). The input layer combines sensory input with a modulatory location input to form sparse representations that correspond to features at specific locations on the object. The output layer receives feedforward inputs from the input layer and converges to a stable pattern representing the object (e.g. a coffee cup). Convergence in the second layer is achieved via two means. One is by integration over time as the sensor moves relative to the object, and the other is via modulatory lateral connections between columns that are simultaneously sensing different locations on the same object (blue arrows in upper layer). Feedback from the output layer to the input layer allows the input layer to predict what feature will be present after the next movement of the sensor. **B**. Pyramidal neurons have three synaptic integration zones, proximal (green), basal (blue), and apical (purple). Although individual synaptic inputs onto basal and apical dendrites have a small impact on the soma, co-activation of a small number of synapses on a dendritic segment can trigger a dendritic spike (*top right*). The HTM neuron model incorporates active dendrites and multiple synaptic integration zones (*bottom*). Patterns recognized on proximal dendrites generate action potentials. Patterns recognized on the basal and apical dendrites depolarize the soma without generating an action potential. Depolarization is a predictive state of the neuron. Our network model relies on these properties and our simulations use HTM neurons. A detailed walkthrough of the algorithm can be found in the supplementary video.

Below we describe our neuron model, the connectivity of layers and columns, and how the sensory and location inputs are combined over time to recognize objects. A more detailed description of the activation and learning rules is available in the Methods section.

#### Neuron model

We use HTM model neurons in the network (Hawkins and Ahmad, 2016). HTM neurons incorporate dendritic properties of pyramidal cells (Spruston, 2008), where proximal, basal, and apical dendritic segments have different functions (**Figure 1B**). Patterns detected on proximal dendrites represent feedforward driving input, and can cause the cell to become active. Patterns recognized on a neuron’s basal and apical dendrites represent modulatory input, and will cause a dendritic spike and depolarize the cell without immediate activation. Depolarized cells fire sooner than, and thereby inhibit, non-depolarized cells that recognize the same feedforward patterns. In the rest of the paper we refer to proximal dendritic inputs as feedforward inputs, and the distal basal and apical dendritic inputs as modulatory inputs. A detailed description of functions of different dendritic integration zones can be found in (Hawkins and Ahmad, 2016).

#### Input layer

The input layer of each cortical column consists of HTM neurons arranged in mini-columns. (Here a mini-column denotes a thin vertical arrangement of neurons (Buxhoeveden, 2002).) In our simulations we typically have 150-250 mini-columns per cortical column, with 16 cells per mini-column (corresponding to 2400 to 4000 cells). The feedforward input of cells in this layer is the sensory input. As in (Hawkins and Ahmad, 2016) cells within a mini-column recognize the same feedforward patterns (Jones, 2000). We map each sensory feature to a sparse set of mini-columns.

The basal modulatory input for cells in the input layer represents the location on an object. During learning, one cell in each active mini-column is chosen to learn the current location signal. During inference, cells that recognize both the modulatory location input and the feedforward driving input will inhibit other cells in the mini-column. In this way, the input layer forms a sparse representation that is unique for a specific sensory feature at a specific location on the object.

#### Output layer

The output layer also contains HTM neurons. The set of active cells in the output layer represents objects. Cells in the output layer receive feedforward driver input from the input layer. During learning, the set of cells representing an object remains active over multiple movements and learns to recognize successive patterns in the input layer. Thus, an object comprises a representation in the output layer, plus an associated set of feature/location representations in the input layer.

The modulatory input to cells in the output layer comes from other output cells representing the same object, both from within the column as well as from neighboring columns via long-range lateral connections. As in the input layer, the modulatory input acts as a bias. Cells with more modulatory input will win and inhibit cells with less modulatory input. Cells representing the same object will positively bias each other. Thus, if a column has feedforward support for objects A and B at time t, and feedforward support for objects B and C at time t+1, the output layer will converge onto the representation for object B at time t+1 due to modulatory input from time t. Similarly, if column 1 has feedforward support for objects A and B, and column 2 has feedforward support for objects B and C, the output layer in both columns will converge onto the representation for object B.

**Figure 2.**
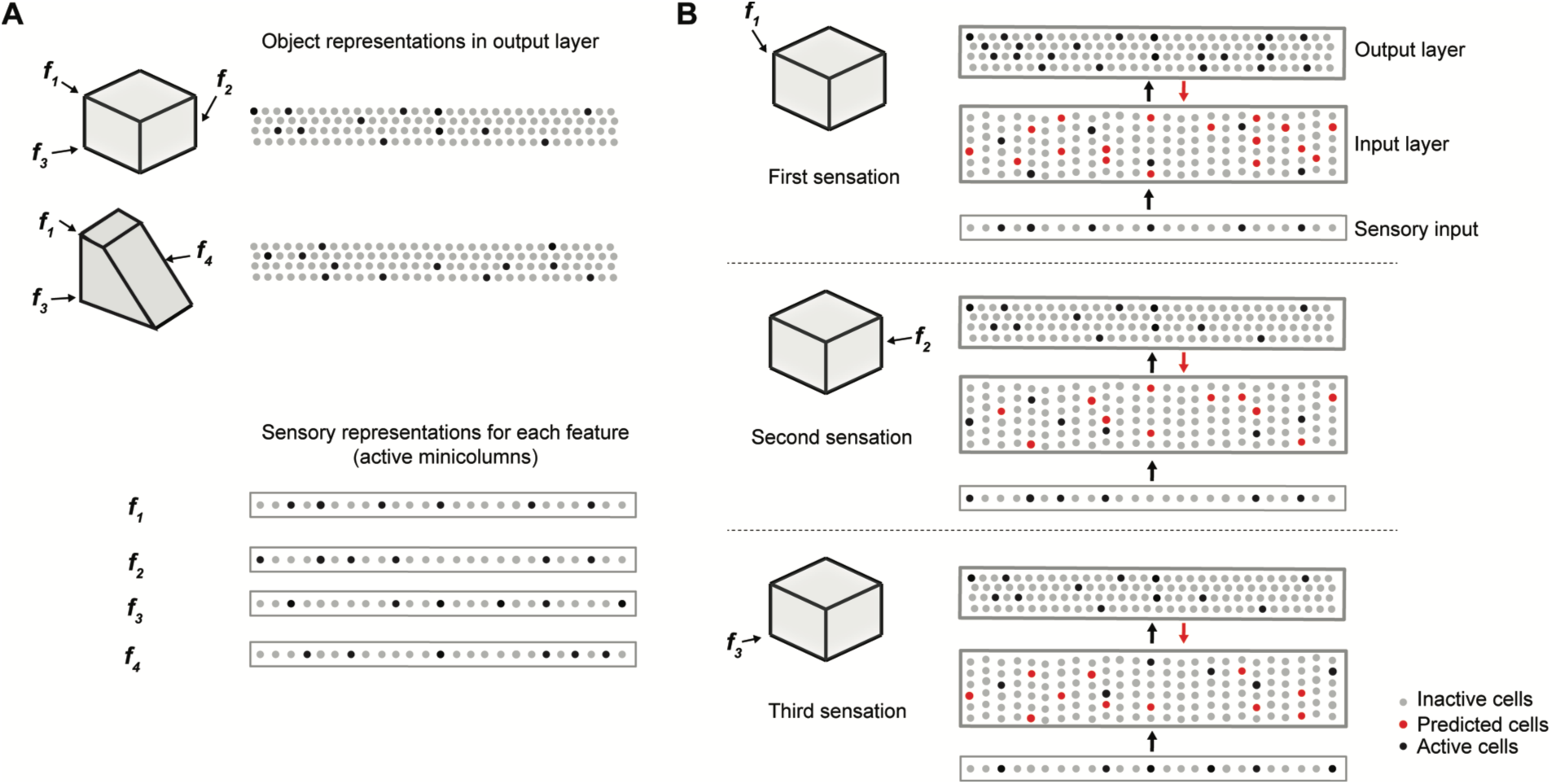
Cellular activations in the input and output layers of a single column during a sequence of touches on an object. **A.** Two objects (cube and wedge). For each object, three feature-location pairs are shown (**f_1_** and **f_3_** are common to both the cube and wedge). The output layer representations associated with each object, and the sensory representations for each feature are shown. **B**. Cellular activations in both layers caused by a sequence of three touches on the cube (in time order from top to bottom). The first touch (at **f_1_**) results in a set of active cells in the input layer (black dots in input layer) corresponding to that feature-location pair. Cells in the output layer receive this representation through their feed-forward connections (black arrow). Since the input is ambiguous, the output layer forms a representation that is the union of both the cube and the wedge (black dots in output layer). Feedback from the output layer to the input layer (red arrow) causes all feature-location pairs associated with both potential objects to become predicted (red dots). The second touch (at **f_2_**) results in a new set of active cells in the input layer. Since **f_2_** is not shared with the wedge, the representation in the output layer is reduced to only the cube. The set of predicted cells in the input layer is also reduced to the feature-location pairs of the cube. The third touch (at **f_3_**) would be ambiguous on its own, however, due to the past sequence of touches and self-reinforcement in the output layer, the representation of the object in the output layer remains unique to the cube. Note the number of cells shown is unrealistically small for illustration clarity.

#### Feedback connections

Neurons in the input layer receive feedback connections from the output layer. Feedback input representing an object, combined with modulatory input representing an anticipated new location due to movement, allows the input layer to more precisely predict the next sensory input. In our model, feedback is an optional component. If included, it improves robustness to sensory noise and ambiguity of location.

### Illustrative example

**Figure 2** illustrates how the two layers of a single cortical column cooperate to disambiguate objects that have shared features, in this case a cube and a wedge. The first sensed feature-location, labeled **f_1_**, is ambiguous, as it could be part of either object. Therefore, the output layer simultaneously invokes a union of representations, one for each object that has that feature at that location. Feedback from the output layer to the input layer puts cells in a predictive state (shown in red). The predicted cells represent the set of all feature-locations consistent with the set of objects active in the output layer. The red cells thus represent the predictions of the network consistent with the sensations up to this point. Upon the second sensation, labeled **f_2_**, only the subset of cells that is consistent with these predictions and the new feature become active. Each subsequent sensation narrows down the set until only a single object is represented in the output layer. A detailed walkthrough of the algorithm can be found in the supplementary video.

### Learning

Learning is based on simple Hebbian-style adaptation : when cells fire, previously active synapses are strengthened and inactive ones are weakened. There are two key differences with most other neural models. First, learning is isolated to individual dendritic segments (Stuart and Häusser, 2001; Losonczy et al., 2008). Second, the model neuron learns by growing and removing synapses from a pool of potential synapses (Chklovskii et al., 2004). We model the growth and removal of synapses by incrementing or decrementing a variable we call “permanence”. The efficacy, or weight, of a synapse is binary based on a threshold of permanence. Thus, how fast the system learns and how long memory is retained can be adjusted independent of the weight of synapses. A complete description of the biological motivation can be found in (Hawkins and Ahmad, 2016). Below we briefly describe how these principles enable the network to learn; the formal learning rules are described in the Materials and Methods section.

**Figure 3.**
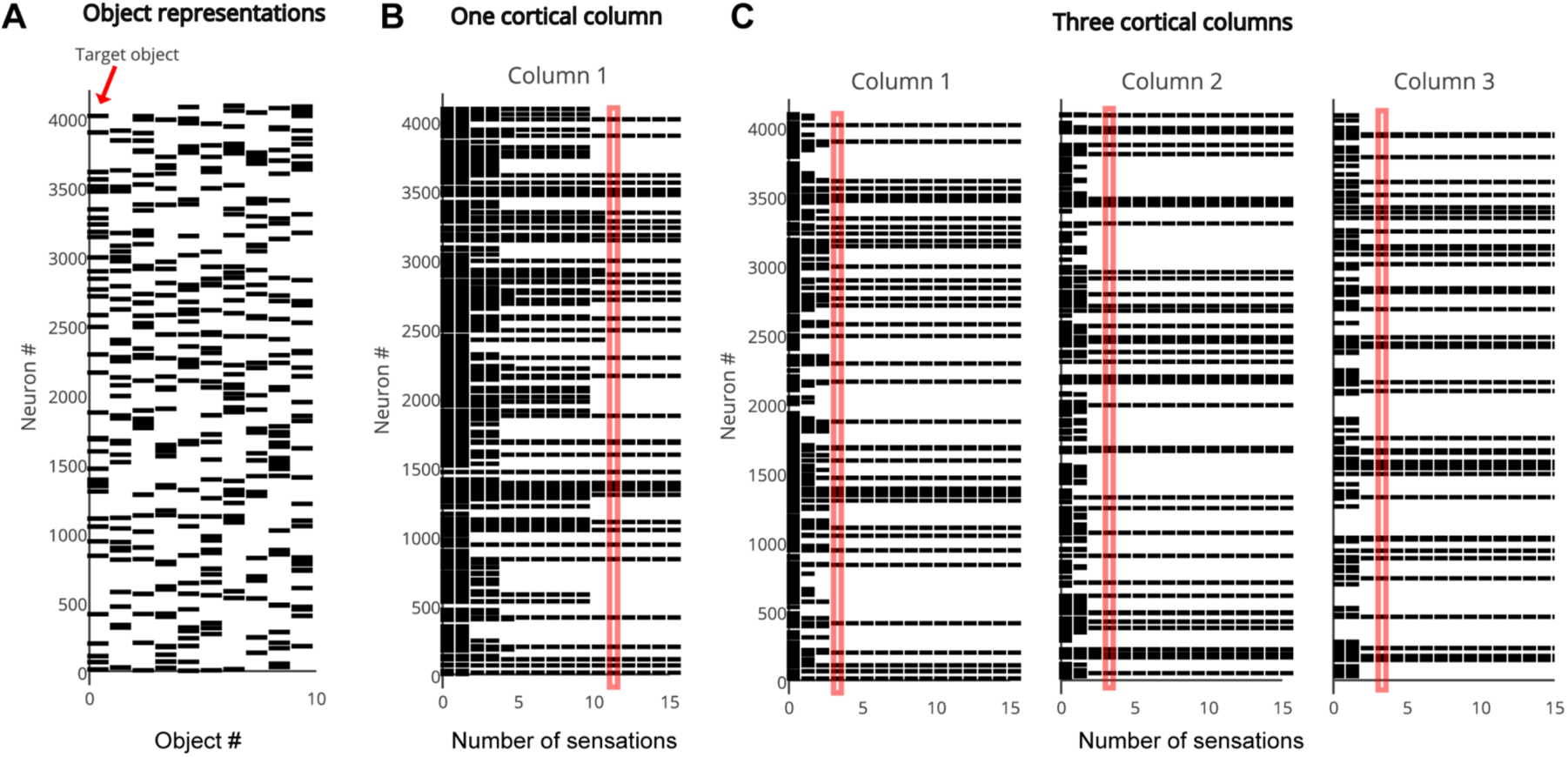
The output layer represents each object by a sparse pattern. We tested the network on the first object. **B**. Activity in the output layer of a single column network as it touches the object. The network converges after 11 sensations (red rectangle). **C**. Activity in the output layer of a three column network as it touches the object. The network converges much faster, after 4 sensations (red rectangle). In both B and C the representation in Column 1 is the same as the target object representation after convergence.

The input layer learns specific feature/location combinations. If the current feature/location combination has not been previously learned (no cell is predicted), then one cell from each active mini-column is chosen as the winner and becomes active. The winning cell is chosen as the cell with the best modulatory input match via random initial conditions. Each winner cell learns by forming and strengthening modulatory connections with the current location input. If the location input is encountered again the corresponding set of cells will be predicted. If the expected sensory feature arrives, the predicted cells will fire first, and the corresponding modulatory inputs will be reinforced. Apical dendrites of the winning cells form connections to active cells in the output layer.

The output layer learns representations corresponding to objects. When the network first encounters a new object, a sparse set of cells in the output layer is selected to represent the new object. These cells remain active while the system senses the object at different locations. Feed forward connections between the changing active cells in the input layer and unchanging active cells in the output layer are continuously reinforced. Thus, each output cell pools over multiple feature/location representations in the input layer. Dendritic segments on cells in the output layer learn by forming lateral modulatory connections to active cells within their own column, and to active cells in nearby columns.

During training, we reset the output layer when switching to a new object. In the brain, there are several ways the equivalent of a reset could occur, including a sufficiently long period of time with no sensation. When a new object is learned we select the object representation based on best match via random initial connectivity.

## SIMULATION RESULTS

In this section, we describe simulation results that illustrate the performance of our network model. The network structure consists of one or more cortical columns, each with two layers, as described earlier (**Figure 1**). In the first set of simulations the input layer of each column consists of 150 mini-columns, with 16 cells per mini-column, for a total of 2400 cells. The output layer of each column consists of 4096 cells, which are not arranged in mini-columns. The output layer contains inter-column and intra-column connections via the distal basal dendrites of each cell. The output layer also projects back to the apical dendrites of the input layer within the same column. All connections are continuously learned and adjusted during the training process.

We trained the network on a library of up to 500 objects (**Figure 2A**). Each object consists of 10 sensory features chosen from a library of 5 to 30 possible features. Each feature is assigned a corresponding location on the object. Note that although each object consists of a unique set of features/locations, any given feature or feature/location is shared across several objects. As such, a single sensation by a single column is insufficient to unambiguously identify an object.

The set of active cells in the output layer represents the objects that are recognized by the network. During inference we say that the network unambiguously recognizes an object when the representation of the output layer overlaps significantly with the representation for correct object and not for any other object. (Complete details of object construction and recognition are described in Materials and Methods).

**Figure 4.**
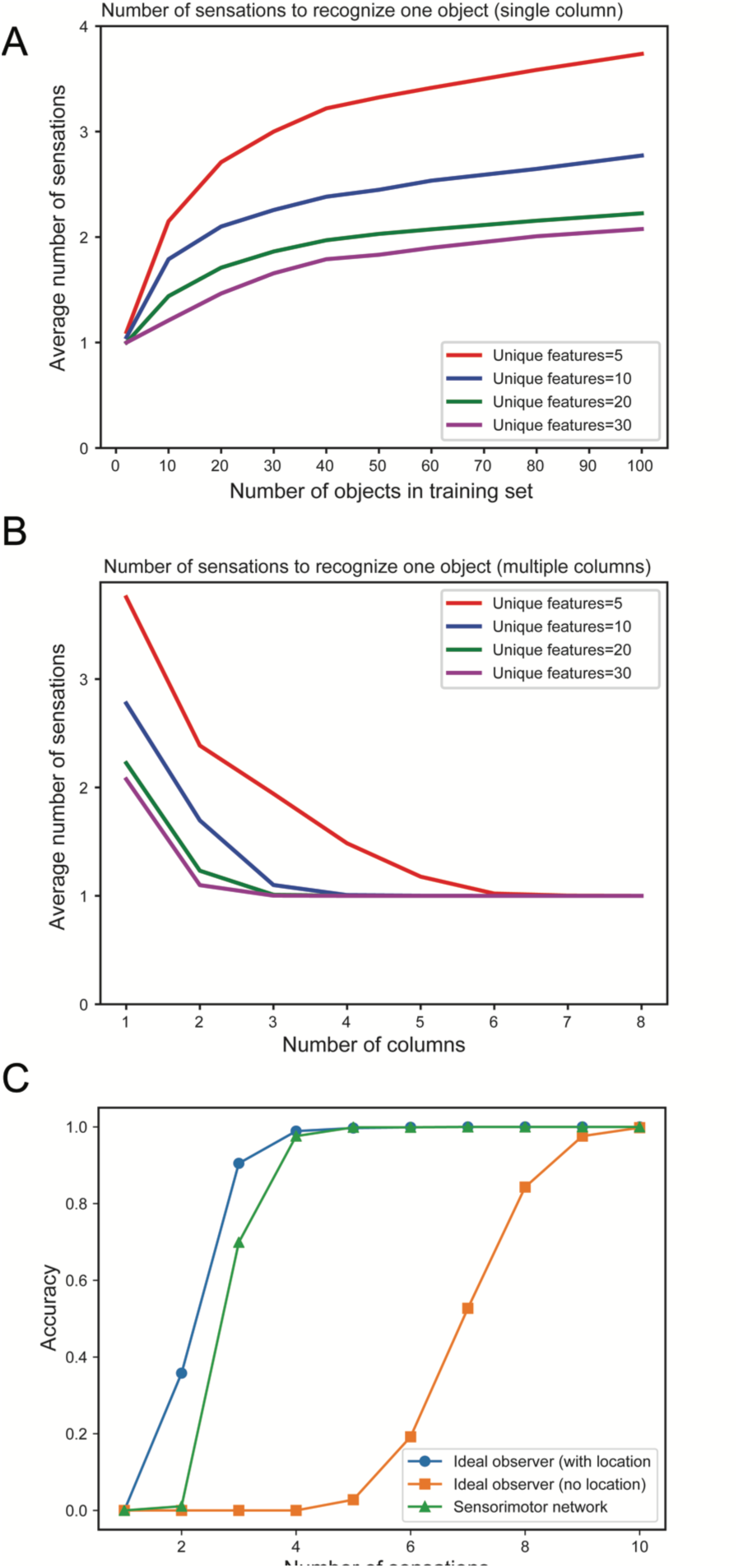
**A**. Mean number of sensations needed to unambiguously recognize an object with a single column network as the set of learned objects increases. We train models on varying numbers of objects, from 1 to 100 and plot the average number of sensations required to unambiguously recognize a single object. The different curves show how convergence varies with the total number of unique features from which objects are constructed. In all cases the network eventually recognizes the object. Recognition requires fewer sensations when the set of features is greater. **B**. Mean number of observations needed to unambiguously recognize an object with multi-column networks as the set of columns increases. We train each network with 100 objects and plot the average number of sensations required to unambiguously recognize an object. The required number of sensations rapidly decreases as the number of columns increases, eventually reaching one. **C.** Fraction of objects that can be unambiguously recognized as a function of number of sensations for an ideal observer model with location (blue), without location (orange) and our one-column sensorimotor network (green).

In the following paragraphs we first describe network convergence, using single and multi-column networks. We then discuss the capacity of the network.

### Network convergence

As discussed earlier, the representation in the output layer is consistent with the recent sequence of sensed features and locations. Multiple output representations will be active simultaneously if the sensed features and locations are not unique to one particular object. The output converges to a single object representation over time as the object is explored via movement. **Figure 3** illustrates the rate of convergence for a one column network and for a three-column network. Multiple columns working together reduces the number of sensations needed for recognition.

In **Figure 4A** we plot the mean number of sensations required to unambiguously recognize an object as a function of the total number of objects in the training set. As expected, the number of sensations required increases with the total number of stored objects. However, in all cases the network eventually correctly recognizes every object. The number of sensations is also dependent on the overall confusion between the set of objects. The more unique the objects, the faster the network can disambiguate them.

**Figure 4B** illustrates the mean number of sensations required to recognize an object as a function of the number of cortical columns in the network. The graph demonstrates the advantage of including multiple columns. The number of sensations required decreases rapidly as the number of columns increases. Thus, although single column networks can recognize the objects, multicolumn networks are much faster. With a sufficient number of columns, the network disambiguates even highly confusing objects with a single sensation. In this experiment, each column receives lateral input from every other column.

In **Figure 4C** we plot the fraction of objects that can be unambiguously recognized (“accuracy”) as a function of the number of sensations. We compare a single column network to an ideal observer model with and without location (see Methods). The performance of our model is close to the ideal observer with locations. It takes many more sensations for the model without locations to recognize the objects, and some objects cannot be distinguished (this is not shown on the graph as we plot the average number of sensations), underscoring the importance of the location signal. We have also shown that multi-column networks perform close to an ideal observer model that similarly observes multiple features per sensation (Supplementary Fig. 9). Taken together, these results show that our biologically derived sensorimotor network operates close to the non-biological ideal model with respect to accuracy and speed of convergence.

### Capacity

In the network model presented here, each cortical column builds predictive models of objects. A key question is, how many objects can a single column represent? Also, does adding more columns impact capacity? In this section we explore the effect of various parameters on the number of objects that can be accurately recognized. We define capacity as the maximum number of objects a network can learn and recognize without confusion. We analyze four different factors that impact capacity : the representational space of the network, the number of mini-columns in the input layer, the number of neurons in the output layer, and the number of cortical columns. In our analysis we used numbers similar to those reported in experimental data. For example, cortical columns vary from 300 μ m to 600 μ m in diameter (Mountcastle, 1997), where the diameter of a mini-column is estimated to be in the range of 30-60 μ m (Buxhoeveden, 2002). For our analysis and simulations we assumed a cortical column contains between 150 and 250 mini-columns.

First, the neural representation must allow the input and output layers to represent large numbers of unique feature/locations and objects. As illustrated in **Figure 2**, both layers use sparse representations. Sparse representations have several attractive mathematical properties that allow robust representation of a very large number of elements (Ahmad and Hawkins, 2016). With a network of 150 mini-columns, 16 cells per mini-column, and 10 simultaneously active mini-columns, we can uniquely represent 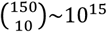 sensory features. Each feature can be represented at 16^10^ unique locations. Similarly, the output layer can represent 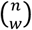unique objects, where *n* is the number of output cells and *w* is the number of active cells at any time. With such large representational spaces, it is extremely unlikely for two feature/location pairs or two object representations to have a significant number of overlapping bits by chance (Supplementary material). Therefore, the number of objects and feature location pairs that can be uniquely represented is not a limiting factor in the capacity of the network.

**Figure 5.**
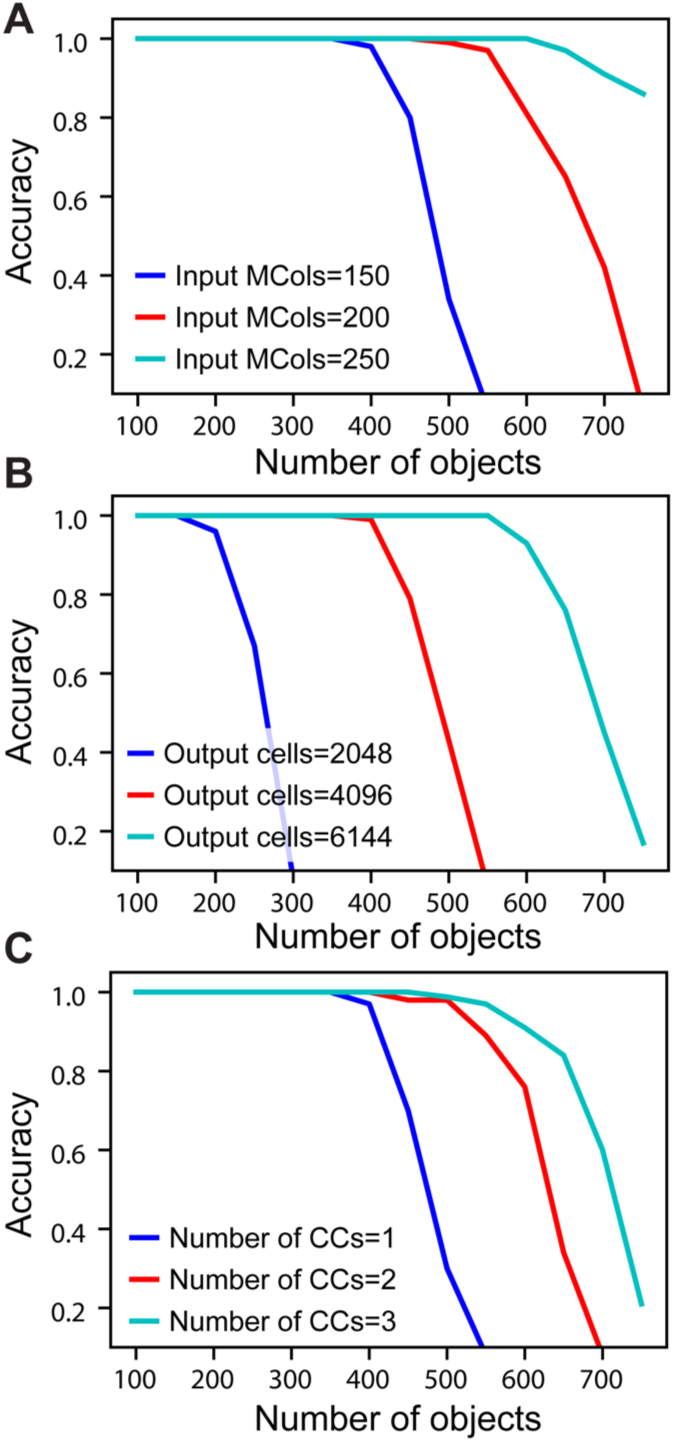
Recognition accuracy is plotted as a function of the number of learned objects. **A**. Network capacity relative to number of mini-columns in the input layer. The number of output cells is kept at 4096 with 40 cells active at any time. **B**. Network capacity relative to number of cells in the output layer. The number of active output cells is kept at 40. The number of mini-columns in the input layer is 150. **C.** Network capacity for one, two, and three cortical columns (CCs). The number of mini-columns in the input layer is 150, and the number of output cells is 4096.

**Figure 6.**
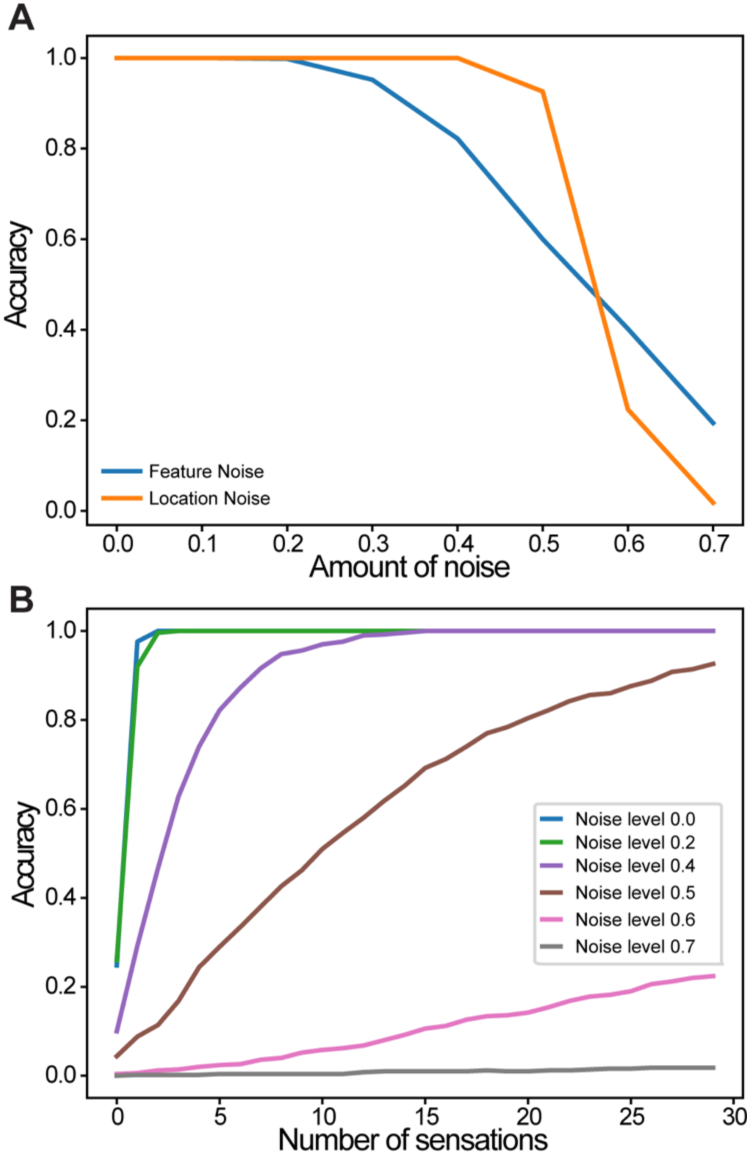
Robustness of a single column network to noise. **A**. Recognition accuracy is plotted as a function of the amount of noise in the sensory input (blue) and in the location input (yellow). **B**. Recognition accuracy as a function of the number of sensations. Colored lines correspond to noise levels in the location input.

As the number of learned objects increases, neurons in the output layer form increasing numbers of connections to neurons in the input layer. If an output neuron connects to too many input neurons, it may be falsely activated by a pattern it was not trained on. Therefore, the capacity of the network is limited by the pooling capacity of the output layer. Mathematical analysis suggests that a single cortical column can store hundreds of objects before reaching this limit (see Supplementary material).

To measure actual network capacity we trained networks with an increasing number of objects and plotted recognition accuracy. For a single cortical column, with 4,096 cells in the output layer and 150 mini-columns in the input layer, the recognition accuracy remains perfect up to 400 objects (**Figure 5A**, blue). The retrieval accuracy drops when the number of learned objects exceeds the capacity of the network.

From the mathematical analysis, we expect the capacity of the network to increase as the size of the input and output layers increase. We again tested our analysis through simulations. With the number of active cells fixed, the capacity increases with the number of mini-columns in the input layer (**Figure 5A**). This is because with more cells in the input layer, the sparsity of activation increases, and it is less likely for an output cell to be falsely activated. The capacity also significantly increases with the number of output cells when the size of the input layer is fixed (**Figure 5B**). This is because the number of feedforward connections per output cell decreases when there are more output cells available. We found that if the size of individual columns is fixed, adding columns can increase capacity (**Figure 5C**). This is because the lateral connections in the output layer can help disambiguate inputs once individual cortical columns hit their capacity limit. However, this effect is limited; the incremental benefit of additional columns decreases rapidly.

The above simulations demonstrate that it is possible for a single cortical column to model and recognize several hundred objects. Capacity is most impacted by the number of cells in the input and output layers. Increasing the number of columns has a marginal effect on capacity. The primary benefit of multiple columns is to dramatically reduce the number of sensations needed to recognize objects. A network with one column is like looking at the world through a straw; it can be done, but slowly and with difficulty.

### Noise robustness

We evaluated robustness of a single column network to noise. After the network learned a set of objects, we added varying amounts of random noise to the sensory and location inputs. The noise affected the active bits in the input without changing its overall sparsity (see Methods). Recognition accuracy after 30 touches is plotted as a function of noise (**Fig. 6A**). There is no impact on the recognition accuracy up to 20% noise in the sensory input and 40% noise in the location input. We also found that the convergence speed was impacted by noise in the location input (**Fig. 6B**). It took more sensations to recognize the object when the location input is noisy.

## MAPPING TO BIOLOGY

Anatomical evidence suggests that the sensorimotor inference model described above exists at least once in each column (layers 4 and 2/3) and perhaps twice (layers 6a and 5). We adopt commonly used terminology to describe these layers. This is a convenience as the connectivity and physiology of cell populations is what matters. Cells we describe as residing in separate layers may actually intermingle in cortical tissue (Guy and Staiger, 2017).

### Layers 4 and 2/3

The primary instance of the model involves layers 4 and 2/3 as illustrated in **Figure 7A**. The following properties evident in L4 and L2/3 match our model. L4 cells receive direct thalamic input from sensory “core” regions (e.g., LGN) (Douglas and Martin, 2004). This input onto proximal dendrites exhibits driver properties (Viaene et al., 2011a). L4 cells do not form long range connections within their layer (Luhmann et al., 1990). L4 cells project to and activate cells in L2/3 (Lohmann and Rörig, 1994; Feldmeyer et al., 2002; Sarid et al., 2007), and receive feedback from L2/3 (Lefort et al., 2009; Markram et al., 2015). L2/3 cells project long distances within their layer (Stettler et al., 2002; Hunt et al., 2011) and are also a major output of cortical columns (Douglas and Martin, 2004; Shipp et al., 2007). It is known that L2/3 activation follows L4 activation (Constantinople and Bruno, 2013).

The model predicts that a representation of location is input to the basal distal dendrites of the input layer. A timing requirement of our model is that the location signal is a predictive signal that must precede the arrival of the sensory input. This is illustrated by the red line in **Figure 7A**. About 45% of L4 synapses come from cells in L6a (Binzegger et al., 2004). The axon terminals were found to show a strong preference for contacting basal dendrites (McGuire et al., 1984) and activation of L6a cells caused weak excitation of L4 cells (Kim et al., 2014). Therefore, we propose that the location representation needed for the upper model comes from L6a.

**Figure 7.**
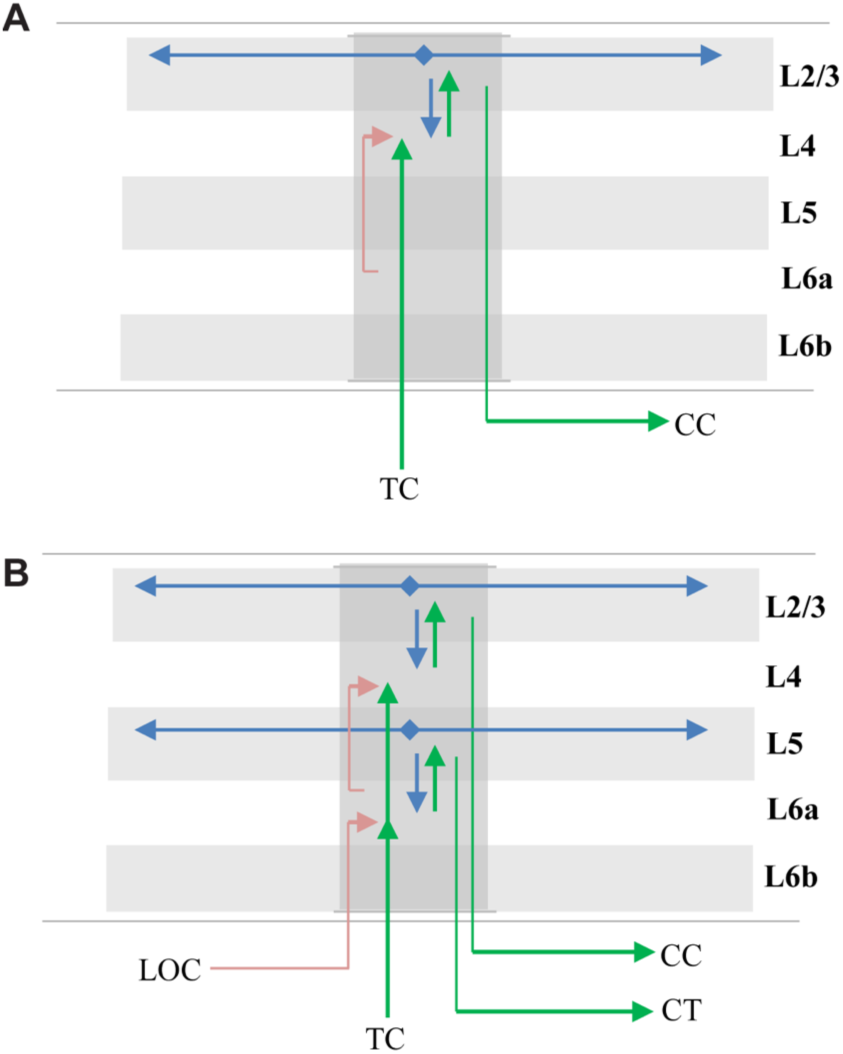
Mapping of sensorimotor inference network onto experimentally observed cortical connections. Arrows represent documented pathways. **A**. First instance of network; L4 is input layer, L2/3 is output layer. Green arrows are feedforward pathway, from thalamo-cortical (TC) relay cells, to L4, to L2/3 cortico-cortical (CC) output cells. Cells in L2/3 also project back to L4 and to adjacent columns (blue arrows); these projections depolarize specific sets of cells that act as predictions (see text). Red arrow is location signal originating in L6a and terminating on basal distal dendrites of L4 cells. **B**. Possible second instance of network; L6a is input layer, L5 is output layer. Both instances of the network receive feedforward input from the same TC axons, thus the two networks run in parallel (Constantinople and Bruno, 2013; Markov et al., 2013). The origin and derivation of the location signal (LOC) is unknown but likely involves local processing as well as input from other regions (see text and Discussion). The output of the upper network makes direct cortical-cortical (CC) connections, whereas the output of the lower network projects to thalamic relay cells before projecting to the next region.

### Layers 6a and 5

Another potential instance of the model is in layers 6a and 5 as illustrated in **Figure 7B**. The following properties evident in L6a and L5 match our model. L6a cells receive direct thalamic input from sensory “core” regions (e.g., LGN) (Thomson, 2010). This input exhibits driver properties and resembles the thalamocortical projections to L4 (Viaene et al., 2011b). L6a cells project to and activate cells in L5 (Thomson, 2010). Recent experimental studies found that the axons of L6 CT neurons densely ramified within layer 5a in both visual and somatosensory cortices of the mouse, and activation of these neurons generated large excitatory postsynaptic potentials (EPSPs) in pyramidal neurons in layer 5a (Kim et al., 2014). L6a cells receive feedback from L5 (Thomson, 2010). L5 cells project long distances within their layer (Schnepel et al., 2015) and L5 cells are also a major output of cortical columns (Douglas and Martin, 2004; Guillery and Sherman, 2011; Sherman and Guillery, 2011). There are three types of pyramidal neurons in L5 (Kim et al., 2015). Here we are referring to only one of them, the larger neurons with thick apical trunks that send an axon branch to relay cells in the thalamus (Ramaswamy and Markram, 2015). However, there is also empirical evidence our model does not map cleanly to L6a and L5. For example, (Constantinople and Bruno, 2013) have shown a sensory stimulus will often cause L5 cells to fire simultaneously or even slightly before L6 cells, which is inconsistent with the model. Therefore, whether L6a and L5 can be interpreted as an instance of the model is unclear.

### Origin of location signal

The derivation of the location representation in L6a is unknown. Part of the answer will involve local processing within the lower layers of the column and part will likely involve long range connections between corresponding regions in “what” and “where” pathways (Thomson, 2010). Parallel “what” and “where” pathways exist in all the major sensory modalities (Ungerleider and Haxby, 1994; Ahveninen et al., 2006). Evidence suggests that regions in “what” pathways form representations that exhibit increasing invariance to translation, rotation or scale and increasing selectivity to sensory features in object centered coordinates (Rust and DiCarlo, 2010). This effect can be interpreted as forming allocentric representations. In contrast, it has been proposed that regions in “where” pathways form representations in egocentric coordinates (Goodale and Milner, 1992). If an egocentric motor behavior is generated in a “where” region, then a copy of the motor command will need to be sent to the corresponding “what” region where it can be converted to a new predicted allocentric location. The conversion is dependent on the current position and orientation of the object relative to the body. It is for this reason we suggest that the origin of the location signal might involve long-range connections between “where” and “what” regions. In the Discussion section we will describe how the location might be generated.

### Physiological evidence

In addition to anatomical support, there are several physiological predictions of the model that are supported by empirical observation. L4 and L6a cells exhibit “simple” receptive fields (RFs) while L2/3 and L5 cells exhibit “complex” RFs (Hubel and Wiesel, 1962; Gilbert, 1977). Key properties of complex cells include RFs influenced by a wider area of sensory input and increased temporal stability (Movshon et al., 1978). L2/3 cells have receptive fields that are twice the size of L4 cells in the primary somatosensory cortex (Chapin, 1986). A distinct group of cells with large and non-oriented receptive fields were found mostly in layer 5 of the visual cortex (Mangini and Pearlman, 1980; Lemmon and Pearlman, 1981). These properties are consistent with, and observed, in the output layer of our model.

The model predicts that cells in a mini-column in the input layer (L4 and L6a) will have nearly identical RFs when presented with an input than cannot be predicted as part of a previously learned object. However, in the context of learned objects, the cells in a mini-column will differentiate. One key differentiation is that individual cells will respond only in specific contexts. This differentiation has been observed in multiple modalities (Vinje and Gallant, 2002; Yen et al., 2006; Martin and Schröder, 2013; Gavornik and Bear, 2014). Our model is also consistent with findings that early sensory areas are biased toward recent perceptual recognition results (St. John-Saaltink et al., 2016).

A particularly relevant version of this phenomenon is “border ownership” (Zhou et al., 2000). Cells which have similar classic receptive fields when presented with isolated edge-like features, diverge and fire uniquely when the feature is part of a larger object. Specifically, the cells fire when the feature is at a particular location on a complex object, a behavior predicted and exhibited by our model. To explain border ownership, researchers have proposed a layer of cells that perform “grouping” of inputs. The grouping cells are stable over time (Craft et al., 2007). The output layer of our model performs this function. “Border ownership” is a form of complex object modeling. It has been observed in both primary and secondary sensory regions (Zhou et al., 2000). We predict that similar properties can be observed in primary and secondary sensory regions for even more complex and three-dimensional objects.

Lee, Carvell, et. al show that enhancement of motor cortex activity facilitates sensory-evoked responses of topographically aligned neurons in primary somatosensory cortex (Lee et al., 2008). Specifically, they found that S1 corticothalamic neurons in whisker/barrel cortex responded more robustly to whisker deflections when motor cortex activity was focally enhanced. This supports the model hypothesis that behaviorally-generated location information projects in a column-by-column fashion to primary sensory regions.

## DISCUSSION

### Relationship with previous models

Due to the development of new experimental techniques, knowledge of the laminar circuitry of the cortex continues to grow (Thomson and Bannister, 2003; Thomson and Lamy, 2007). It is now possible to reconstruct and simulate the circuitry in an entire cortical column (Markram et al., 2015). Over the years, numerous efforts have been undertaken to develop models of cortical columns. Many cortical column models aim to explain neurophysiological properties of the cortex. For example, based on their studies on the cat visual cortex, (Douglas and Martin, 1991) provided one of the first canonical microcircuit models of a cortical column. This model explains intracellular responses to pulsed visual stimulations and has remained highly influential (Douglas and Martin, 2004). (Hill and Tononi, 2004) constructed a large-scale model of point neurons that are organized in a repeating columnar structure to explain the difference of brain states during sleep and wakefulness. (Traub et al., 2004) developed a single-column network model based on multi-compartmental biophysical models to explain oscillatory, epileptic and sleeplike phenomena. (Reimann et al., 2013) showed that the neocortical local field potentials can be explained by a cortical column model composed of >12,000 reconstructed multi-compartmental neurons.

Although these models provided important insights on the origin of neurophysiological signals, there are relatively few models proposing the functional roles of layers and columns. (Bastos et al., 2012) discussed the correspondence between the micro-circuitry of the cortical column and the connectivity implied by predictive coding. This study used a coarse microcircuit model based on the work of (Douglas and Martin, 2004) and lacked recent experimental evidence and detailed connectivity patterns across columns.

(Raizada and Grossberg, 2003) described the LAMINART model to explain how attention might be implemented in the visual cortex. This study highlighted the anatomical connections of the L4-L2/3 network and proposed that perceptual grouping relies on long-range lateral connections in L2/3. This is consistent with our proposal of the stable object representation in L2/3. A recent theory of optimal context integration proposes that long-range lateral connections are used to optimally integrate information from the surround (Iyer and Mihalas, 2017). The structure of their model is broadly consistent with the theories presented here, and provides a possible mathematical basis for further analysis.

### The benefit of cortical columns

Our research has been guided by Mountcastle’s definition of a cortical column (Mountcastle, 1978, 1997), as a structure ‘formed by many mini-columns bound together by short-range horizontal connections ’. The concept plays an essential role in the theory presented in this paper. Part of our theory is that each repetitive unit, or “column”, of sensory cortex can learn complete objects by locally integrating sensory and location data over time. In addition, we have proposed that multiple cortical columns greatly speed up inference and recognition time by integrating information in parallel across dispersed sensory areas.

An open issue is the exact anatomical organization of columns. We have chosen to describe a model of columns with discrete inter-column boundaries. This type of well-defined structure is most clear in the rat barrel cortex (Lubke et al., 2000; Bureau et al., 2004; Feldmeyer et al., 2013) but Mountcastle and others have pointed out that although there are occasional discontinuities in physiological and anatomical properties, there is a diverse range of structures and the more general rule is continuity (Mountcastle, 1978; Horton and Adams, 2005; Rockland, 2010).

Mountcastle’s concept of a repetitive functional unit, whether continuous or discrete, is useful to understand the principles of cortical function. Our model assigns a computational benefit to columns, that of integrating discontinuous information in parallel across disparate areas. This basic capability is independent of any specific type of column (such as hypercolumns or ocular dominance columns), and independent of discrete or continuous structures. The key requirement is that each column models a different subset of sensory space and is exposed to different parts of the world as sensors move.

### Generating the location signal

A key prediction of our model is the presence of a location signal in each column of a cortical region. We deduced the need for this signal based on the observation that cortical regions predict new sensory inputs due to movement (Duhamel et al., 1992; Nakamura and Colby, 2002; Li and DiCarlo, 2008). To predict the next sensory input, a patch of neocortex needs to know where a sensor will be on a sensed object after a movement is completed. The prediction of location must be done separately for each part of a sensor array. For example, for the brain to predict what each finger will feel on a given object, it has to predict a separate allocentric location for each finger. There are dozens of semi-independent areas of sensation on each hand, each of which can sense a different location and feature on an object. Thus, the allocentric location signals must be computed in a part of the brain where somatic topology is similarly granular. For touch, this suggests the derivation of allocentric location is occurring in each column throughout primary regions such as S1 and S2. The same argument holds for primary visual regions, as each patch of the retina observes different parts of objects.

Although we don’t know how the location signal is generated, we can list some theoretically-derived requirements. A column needs to know its current location on an object, but it also needs to predict what its new location will be after a movement is completed. To translate an egocentric motor signal into a predicted allocentric location, a column must also know the orientation of the object relative to the body part doing the moving. This can be expressed in the pseudo-equation [current location + orientation of object + movement ⇒ predicted new location]. This is a complicated task for neurons to perform. Fortunately, it is highly analogous to what grid cells do. Grid cells are a proof that neurons can perform these types of transformations, and they suggest specific mechanisms that might be deployed in cortical columns.

1. Grid cells in the entorhinal cortex (Hafting et al., 2005; Moser et al., 2008) encode the location of an *animal’s body* relative to an external *environment*. A sensory cortical column needs to encode the location of a *part of the animal’s body* (a sensory patch) relative to an external *object*.
2. Grid cells use path integration to predict a new location due to movement (Kropff et al., 2015). A column must also use path integration to predict a new location due to movement.
3. To predict a new location, grid cells combine current location, with movement, with head direction cells (Moser et al., 2014). Head direction cells represent the “orientation” of the “animal” relative to an external environment. Columns need a representation of the “orientation” of a “sensory patch” relative to an external object.
4. The representation of space using grid cells is dimensionless. The dimensionality of the space they represent is defined by the tiling of grid cells, combined with how the tiling maps to behavior. Similarly, our model uses representations of location that are dimensionless.

These analogs, plus the fact that grid cells are phylogenetically older than the neocortex, lead us to hypothesize that the cellular mechanisms used by grid cells were preserved and replicated in the sub-granular layers of each cortical column. It is not clear if a column needs neurons that are analogous to place cells (Moser et al., 2015). Place cells are believed to associate a location (derived from grid cells) with features and events. They are believed to be important for episodic memory. Presently, we don’t see an analogous requirement in cortical columns.

Today we have no direct empirical evidence to support the hypothesis of grid-cell like functionality in each cortical column. We have only indirect evidence. For example, to compute location, cortical columns must receive dynamically updated inputs regarding body pose. There is now significant evidence that cells in numerous cortical areas, including sensory regions, are modulated by body movement and position. Primary visual and auditory regions contain neurons that are modulated by eye position (Trotter and Celebrini, 1999; Werner-Reiss et al., 2003) as do areas MT, MST, and V4 (Bremmer, 2000; DeSouza et al., 2002). Cells in frontal eye fields (FEF) respond to auditory stimuli in an eye-centered frame of reference (Russo and Bruce, 1994). Posterior parietal cortex (PPC) represents multiple frames of reference including head-centered (Andersen et al., 1993) and body-centered (Duhamel et al., 1992; Brotchie et al., 1995, 2003; Bolognini and Maravita, 2007) representations. Motor areas also contain a diverse range of reference frames, from representations of external space independent of body pose to representations of specific groups of muscles (Graziano and Gross, 1998; Kakei et al., 2003). Many of these representations are granular, specific to particular body areas, and multisensory, implying numerous transformations are occurring in parallel (Graziano et al., 1997; Graziano and Gross, 1998; Rizzolatti et al., 2014). Some models have shown that the above information can be used to perform coordinate transformations (Zipser and Andersen, 1988; Pouget and Snyder, 2000).

Determining how columns derive the allocentric location signal is a current focus of our research.

### Role of Inhibitory Neurons

There are several aspects of our model that require inhibition. In the input layer, neurons in mini-columns mutually inhibit each other. Specifically, neurons that are partially depolarized (in the predictive state) generate a first action potential slightly before cells that are not partially depolarized. Cells that spike first prevent other nearby cells from firing. This requires a very fast, winner-take-all type of inhibition among nearby cells, and suggests that such fast inhibitory neurons contain stimulus-related information, which is consistent with recent experiment findings (Reyes-Puerta et al., 2015). Simulations of the timing requirement for this inhibition can be found in (Billaudelle and Ahmad, 2015). Activations in the output layer do not require very fast inhibition. Instead, a broad inhibition within the layer is needed to maintain the sparsity of activation patterns. Experiment evidence for both fast and broad inhibition have been reported in the literature (Helmstaedter et al., 2009; Meyer et al., 2011).

Our simulations do not model inhibitory neurons as individual cells. The functions of inhibitory neurons are encoded in the activation rules of the model. A more detailed mapping to specific inhibitory neuron types is an area for future research.

### Hierarchy

The neocortex processes sensory input in a series of hierarchically arranged regions. As input ascends from region to region, cells respond to larger areas of the sensory array and to more complex features. A common assumption is that complete objects can only be recognized at a level in the hierarchy where cells respond to input over the entire sensory array.

Our model proposes an alternate view. All cortical columns, even columns in primary sensory regions, are capable of learning representations of complete objects. However, our network model is limited by the spatial extent of the horizontal connections in the output layer. Therefore, hierarchy is still required in many situations. For example, say we present an image of a printed letter on the retina. If the letter occupies a small part of the retina, then columns in V1 could recognize the letter. If, however, the letter is expanded to occupy a large part of the retina, then columns in V1 would no longer be able to recognize the letter because the features that define the letter are too far apart to be integrated by the horizontal connections in L2/3. In this case, a converging input onto a higher cortical region would be required to recognize the letter. Thus the cortex learns multiple models of objects, both within a region and across hierarchical levels.

What would occur if multiple objects were being sensed at the same time? In our model, one part of a sensory array could be sensing one object and another part of the sensory array could be sensing a different object. Difficulty would arise if the sensations from two or more objects were overlaid or interspersed on a region, such as, if your index and ring finger touched one object while your thumb and middle finger touched another object. In these situations, we suspect the system would settle on one interpretation or the other.

Sensory information is processed in parallel pathways, sometimes referred to as “what” and “where” pathways. We propose that our object recognition model exists in “what” regions, which are associated with the ability to recognize objects. How might we interpret “where” pathways in light of our model? First, the anatomy in the two pathways is similar. This suggests that “what” and “where” regions perform similar operations, but achieve different results by processing different types of data. For example, our network might learn models of ego-centric space if the location signal represented ego-centric locations. Second, we suspect that bi-directional connections between what and where regions are required for converting ego-centric motor behaviors into allocentric locations. We are currently exploring these ideas.

### Vision, audition, and beyond

We described our model using somatic sensation. Does it apply to other sensory modalities? We believe it does. Consider vision. Vision and touch are both based on an array of receptors topologically mapped to an array of cortical columns. The retina is not like a camera. The blind spot and blood vessels prevent all parts of an object from being sensed simultaneously, and the density of receptors in the retina is not uniform. Similarly, the skin cannot sense all parts of an object at once, and the distribution of somatic receptors is not uniform. Our model is indifferent to discontinuities and non-uniformities. Both the skin and retina move, exposing cortical columns to different parts of sensed objects over time. The methods for determining the allocentric location signal for touch and vision would differ somewhat. Somatic sensation has access to richer proprioceptive inputs, whereas vision has access to other clues such as ocular disparity. Aside from differences in how allocentric location is determined, our model is indifferent to the underlying sensory modality. Indeed, columns receiving visual input could be interspersed with columns receiving somatic input, and the long-range intercolumn connections in our model would unite these into a single object representation.

Similar parallels can be made for audition. Perhaps the more powerful observation is that the anatomy supporting our model exists in most, if not all, cortical regions. This suggests that no matter what kind of information a region is processing, its feedforward input is interpreted in the context of a location. This would apply to high-level concepts as well as low-level sensory data. This hints at why it is easier to memorize a list of items when they are mentally associated with physical locations, and why we often use mental imagery to convey abstract concepts.

### Testable predictions

A number of experimentally testable predictions follow from this theory.

1. The theory predicts that sensory regions will contain cells that are stable over movements of a sensor while sensing a familiar object.
2. The set of stable cells will be both sparse and specific to object identity. The cells that are stable for a given object will in general have very low overlap with those that are stable for a completely different object.
3. Layers 2/3 of cortical columns will be able to independently learn and model complete objects. We expect that the complexity of the objects a column can model will be related to the extent of long-range lateral connections.
4. Activity within the output layer of each cortical column (layers 2/3) will become sparser as more evidence is accumulated for an object. Activity in the output layer will be denser for ambiguous objects. These effects will only be seen when the animal is freely observing familiar objects.
5. These output layers will form stable representations. In general, their activity will be more stable than layers without long-range connections.
6. Activity within the output layers will converge on a stable representation slower with long-range lateral connections disabled, or with input to adjacent columns disabled.
7. The theory provides an algorithmic explanation for border ownership cells (Zhou et al., 2000). In general each region will contain cells tuned to the location of features in the object’s reference frame. We expect to see these representations in layer 4.

### Summary

Our research has focused on how the brain makes predictions of sensory inputs. Starting with the premise that all sensory regions make predictions of their constantly changing input, we deduced that each small area in a sensory region must have access to a location signal that represents where on an object the column is sensing. Building on this idea, we deduced the probable function of several cellular layers and are beginning to understand what cortical columns in their entirety might be doing. Although there are many things we don’t understand, the big picture is increasingly clear. We believe each cortical column learns a model of “its” world, of what it can sense. A single column learns the structure of many objects and the behaviors that can be applied to those objects. Through intra-laminar and long-range cortical-cortical connections, columns that are sensing the same object can resolve ambiguity.

In 1978 Vernon Mountcastle reasoned that since the complex anatomy of cortical columns is similar in all of the neocortex, then all areas of the neocortex must be performing a similar function (Mountcastle, 1978). His hypothesis remains controversial partly because we haven’t been able to identify what functions a cortical column performs, and partly because it has been hard to imagine what single complex function is applicable to all sensory and cognitive processes.

The model of a cortical column presented in this paper is described in terms of a sensory regions and sensory processing, but the circuitry underlying our model exists in all cortical regions. Thus, if Mountcastle’s conjecture is correct, even high-level cognitive functions, such as mathematics, language, and science would be implemented in this framework. It suggests that even abstract knowledge is stored in relation to some form of “location” and that much of what we consider to be “thought” is implemented by inference and behavior generating mechanisms originally evolved to move and infer with fingers and eyes.

## MATERIALS AND METHODS

Here we formally describe the activation and learning rules for the HTM sensorimotor inference network. We use a modified version of the HTM neuron model (Hawkins and Ahmad, 2016) in the network. There are three basic aspects of the algorithm : initialization, computing cell states, and learning. These steps are described along with implementation and simulation details.

### Notation

Let *N*^in^ represent the number of mini-columns in the input layer, *M* the number of cells per mini-column in the input layer, *N*^out^the number of cells in the output layer and *N*^c^ the number of cortical columns. The number of cells in the input layer and output layer is *MN*^in^ and *N*^out^ respectively for each cortical column. Each input cell receives both the sensory input and a contextual input that corresponds to the location signal. The location signal is a *N*^ext^ dimensional sparse vector ***L***.

Each cell can be in one of three states : active, predictive, or inactive. We use *M* × *N*^in^ binary matrices **A**^in^ and Π^in^ to denote activation state and predictive state of input cells and use the *N*^out^ dimensional binary vector **A**^out^ to denote the activation state of the output cells in a cortical column. The concatenated output of all cortical columns is represented as a N^out^ N^column^dimensional binary vector 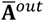 At any point in time there are only a small number of cells active, so these are generally very sparse.

Each cell maintains a single proximal dendritic segment and a set of basal distal dendritic segments (denoted as basal below). Proximal segments contain feedforward connections to that cell. Basal segments represent contextual input. The contextual input acts as a tiebreaker and biases the cell to win. The contextual input to a cell in the input layer is a vector representing the external location signal ***L***. The contextual input to a cell in the output layer comes from other output cells in the same or different cortical columns.

For each dendritic segment, we maintain a set of “potential” synapses between the dendritic segment and other cells that could potentially form a synapse with it (Chklovskii et al., 2004; Hawkins and Ahmad, 2016). Learning is modeled by the growth of new synapses from this set of potential synapses. A “permanence” value is assigned to each potential synapse and represents the growth of the synapse. Potential synapses are represented by permanence values greater than zero. A permanence value close to zero represents an unconnected synapse that is not fully grown. A permanence value greater than the connection threshold represents a connected synapse. Learning occurs by incrementing or decrementing permanence values.

We denote the synaptic permanences of the *d* th dendritic segment of the *i* th input cell in the *j* th mini-column as a *N*^ext^ × 1 vector **D**^ijd,in^. Similarly, the permanences of the *d*th dendritic segment of the *i* th output cell is the *N*^out^*N*^c^ × 1 dimensional vector **D**^id,out^.

Output neurons receive feedforward connections from input neurons within the same cortical column. We denote these connections with a *M* × *N*^in^ × *N*^out^ tensor **F**, where *f*_ijk_ represents the permanence of the synapse between the *i* th input cell in the *j*th mini-column and the *k*th output cell.

For **D** and **F**, we will use a dot (e.g. 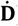) to denote the binary vector representing the subset of potential synapses on a segment (i.e. permanence value above 0). We use a tilde (e.g. 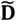) to denote the binary vector representing the subset of connected synapses (i.e. permanence value above connection threshold).

### Initialization

Each dendritic segment is initialized to contain a random set of potential synapses. **D**^ijd,in^ is initialized to contain a random set of potential synapses chosen from the location input. Segments in **D**^id.out^ are initialized to contain a random set of potential synapses to other output cells. These can include cells from the same cortical column. We enforce the constraint that a given segment only contains synapses from a single column. In all cases the permanence values of potential synapses are chosen randomly : initially some are connected (above threshold) and some are unconnected.

### Computing cell states

A cell in the input layer is predicted if any of its basal distal segments have sufficient activity:

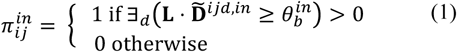

where 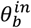 is the activation threshold of the basal distal dendrite of an input cell.

For the input layer, all the cells in a mini-column share the same feedforward receptive fields. Following (Hawkins and Ahmad, 2016) we assume that an inhibitory process selects a set of *s* mini-columns that best match the current feedforward input pattern. We denote this winner set as **W**^in^. The set of active input layer cells is calculated as follows:

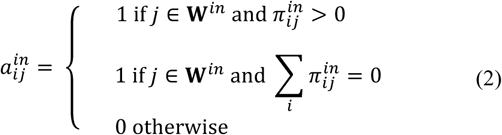

The first conditional states that predicted cells in a winning mini-column becoming winners and become active. If no cell in a mini-column is predicted, all cells in that mini-column become active (second conditional).

To determine activity in the output layer we calculate the feedforward and lateral input to each cell. Cells with enough feedforward overlap with the input layer, and the most lateral support from the previous time step become active. The feedforward overlap to the *k*th output cell is.:

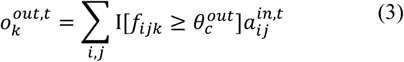

The set of output cells with enough feedforward input is computed as:

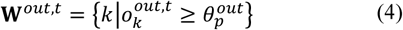

where 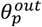 is a threshold. We then select the active cells using the number of active basal segments as a sorting function:

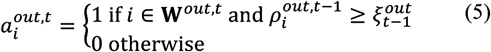

where 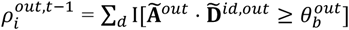 represents the number of active basal segments in the previous time step, and the *s*th highest number of active basal segments is denoted as 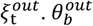 is the activation threshold of the basal distal dendrite of an output cell. *I* [ ] is the indicator function, and *s* is the minimum desired number of active neurons. If the number of cells with lateral support is less than *s* in a cortical column, 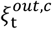 would be zero and all cells with enough feedforward input will become active. Note that we used a modified version of the original HTM neuron model in the output layer by considering the effect of multiple active basal segments.

### Learning in the input layer

In the input layer, basal segments represent predictions. At any point only segments that match its contextual input are modified. If a cell was predicted (Eq. (1)) and becomes active, the corresponding basal segments are selected for learning. If no cell in an active mini-column was predicted, we select a winning cell as the cell with the best basal input match via random initial conditions.

For selected segments, we decrease the permanence of inactive synapses by a small value *p*^-^ and increase the permanence of active synapses by a larger value *p*^+^

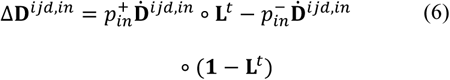

where ∘ represents element-wise multiplication. Incorrect predictions are negatively punished. If a basal dendritic segment on a cell becomes active and the cell subsequently does not become active, we slightly decrement the permanences of active synapses on the corresponding segments. Note that in Eq. 6, learning is applied to all potential synapses (denoted by 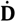).

Learning in the output layer : When learning a new object a sparse set of cells in the output layer is selected to represent the new object. These cells remain active while the system senses the object at different locations. Thus, each output cell pools over multiple feature/location representations in the input layer.

For each sensation, proximal synapses are learned by increasing the permanence of active synapses by 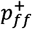, and decreasing the permanence of inactive synapses by 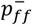:

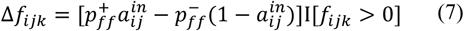

Basal segments of active output cells are learned using a rule similar to (7):

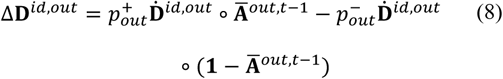

### Feedback

Feedback from the output layer to the input layer is used as an additional modulatory input to fine tune which cells in a winning mini-column become active. Cells in the input layer maintain a set of apical segments similar to the set of basal segments. If a cell has apical support (i.e. an active apical segment), we use a slightly lower value of 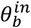 to calculate 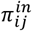 In addition if multiple cells in a mini-column are predicted, only cells with feedback become active. These rules make the set of active cells more precise with respect to the current representation in the output layer. Apical segments on winning cells in the input layer are learned using exactly the same rules as basal segments.

### Simulation details

To generate our convergence and capacity results we generated a large number of objects. Each object consists of a number of sensory features, with each feature assigned to a corresponding location. We encode each location as a 2400-dimensional sparse binary vector with 10 random bits active. Each sensory feature is similarly encoded by a vector with 10 random bits active. The length of the sensory feature vector is the same as the number of mini-columns of the input layer *N*^in^ The input layer contains 150 mini-columns and 16 cells per mini-column, with 10 mini-columns active at any time. The activation threshold of basal distal dendrite of input neuron is 6. The output layer contains 4096 cells and the minimum number of active output cells is 40. The activation threshold is 3 for proximal dendrites and 18 for basal dendrites for output neurons.

During training, the network learns each object in random order. For each object, the network senses each feature three times. The activation pattern in the output layer is saved for each object to calculate retrieval accuracy. During testing, we allow the network to sense each object at *K* locations. After each sensation, we classify the activity pattern in the output layer. We say that an object is correctly classified if, for each cortical column, the overlap between the output layer and the stored representation for the correct object is above a threshold, and the overlaps with the stored representation for all other objects are below that threshold. We use a threshold of 30.

For the network convergence experiment (Figure **4-5**), each object consists of 10 sensory features chosen from a library of 5 to 30 possible features. The number of sensations during testing is 20. For the capacity experiment, each object consists of 10 sensory features chosen from a large library of 5000 possible features. The number of sensations during testing is 3.

Finally, we make some simplifying assumptions that greatly speed up simulation time for larger networks. Instead of explicitly initializing a complete set of synapses across every segment and every cell, we greedily create segments on a random cell and initialize potential synapses on that segment by sampling from currently active cells. This happens only when there is no match to any existing segment.

For the noise robustness experiment (Figure. 6), we added random noise to the sensory input and the location input. For each input, we randomly flip a fraction of the active input bits to inactive, and flip the corresponding number of inactive input bits to active. This procedure randomizes inputs while maintaining constant input sparsity. The noise level denotes the fraction of active input bits that are changed for each input. We varied the amount of noise between 0 and 0.7.

We constructed an ideal observer model to estimate the theoretical upper limit for model performance (Fig. 4C, Supplementary Fig. 9). During learning, the ideal observer model memorizes a list of (feature, location) pairs for each object. During inference, the ideal observer model stores the sequence of observed (feature, location) pairs and calculates the overlap between all the observed pairs and the memorized list of pairs for each object. The predicted object is the object that has the most overlap with all the observed sensations. To compare the ideal observer with a multi-column network with N columns, we provide it with N randomly chosen observations per sensation. Performance of the ideal observer model represents the best one can do given all the sensations up to the current time. We also used the same framework to create a model that only uses sensory features, but no location signals (used in Fig. 4C).

## ACKNOWLEDGEMENTS

We thank Jeff Gavornik for his thoughtful comments and suggestions. We thank the reviewers for their detailed comments, which have helped to improve the paper significantly. We also thank Marcus Lewis, Nathanael Romano, and numerous other collaborators at Numenta over the years for many discussions.

